# SQST-1/p62-regulated SKN-1/Nrf mediates a phagocytic stress response via transcriptional activation of *lyst-1*/LYST

**DOI:** 10.1101/2025.01.08.631988

**Authors:** Aladin Elkhalil, Alec Whited, Piya Ghose

## Abstract

Cells may be intrinsically fated to die to sculpt tissues during development or to maintain homeostasis. Cells can also die in response to various stressors, injury or pathological conditions. Additionally, cells of the metazoan body are often highly specialized with distinct domains that differ both structurally and with respect to their neighbors. Specialized cells can also die, as in normal brain development or pathological states and their different regions may be eliminated via different programs. Clearance of different types of cell debris must be performed quickly and efficiently to prevent autoimmunity and secondary necrosis of neighboring cells. All cells, including those programmed to die, may be subject to various stressors. Some largely unexplored questions include whether predestined cell elimination during development could be altered by stress, if adaptive stress responses exist and if polarized cells may need compartment-specific stress-adaptive programs. We leveraged Compartmentalized Cell Elimination (CCE) in the nematode *C. elegans* to explore these questions. CCE is a developmental cell death program whereby three segments of two embryonic polarized cell types are eliminated differently. We have previously employed this *in vivo* genetic system to uncover a cell compartment-specific, cell non-autonomous clearance function of the fusogen EFF-1 in phagosome closure during corpse internalization. Here, we introduce an adaptive response that serves to aid developmental phagocytosis as a part of CCE during stress. We employ a combination of forward and reverse genetics, CRISPR/Cas9 gene editing, stress response assays and advanced fluorescence microscopy. Specifically, we report that, under heat stress, the selective autophagy receptor SQST-1/p62 promotes the nuclear translocation of the oxidative stress-related transcription factor SKN-1/Nrf. This in turn allows SKN-1/Nrf to transcribe the lysosomal trafficking associated gene *lyst-1*/LYST which subsequently promotes the phagocytic resolution of the developmentally-killed internalized cell even under stress conditions.

**Author Summary:** During development, cells can have many fates, one of which is to deliberately die. If a cell’s inherent ability to die is lost, unwanted cells remain, which can lead to pathologies such as abnormal brain development or cancer. Dead cell remains must also be fully and efficiently cleared away by being ingested and digested by other cells, to avoid autoimmunity. Cells that are destined to die, like any cell, can be subject to stress, which can change cell behavior. Moreover, cells fated to die often have highly intricate shapes, such as nerve cells in the brain, and their removal may entail different strategies for different regions of the cell. In this study, we have used the pre-destined “3-in-1” death of a structurally-complex cell in the roundworm *C. elegans* as a platform to describe the genetics behind how one cell bolsters its inherent ability to consume an area of another dying cell by mounting a response to environmental stress. Specifically, we report, to our knowledge for the first time, that a well-known stress-protective protein helps turns on a gene that helps ensure that ingested parts of dead cells are fully digested and removed.

## Introduction

Programmed cell elimination is an important feature of both normal development and homeostasis [1–3] and entails both cell killing and clearance. Apoptosis is the best characterized form of developmentally programmed cell death marked by defined features and genetics [4, 5]. Several other forms of non-apoptotic or non-canonical regulated cell death programs have been described in recent years [6–10]. Dying cells must subsequently be cleared efficiently via phagocytosis to prevent secondary necrosis and autoimmune consequences. During phagocytosis, cell corpses and debris are internalized following their recognition by phagocytes resulting in the formation of corpse-bearing vesicles called phagosomes that undergo a series of maturation steps [11–15]. Phagosome maturation entails the dead cell cargo-containing vesicle becoming sequentially acidified and subsequently fusing to lysosomes and the resolution of the cargo [16]. Lysosomes are membrane-surrounded acidic organelles consisting of hydrolases, membrane proteins, and numerous accessory proteins. They carry digestive enzymes and are trafficked to the phagosome vesicle allowing for ultimate digestion and resolution of the corpse/debris contained in it. While phagosome maturation has been extensively characterized [11–15], there are still poorly understood factors associated.

All cells are subject to exposure to stress and an integral part of cellular physiology is the ability to adapt and restore homeostasis to ensure normal fate and function. Cells can encounter a myriad of stressors in their lifetime including heat stress, oxidative stress, UV stress, and pathogenic stress, which they combat by mounting appropriate stress responses [17–25], which have some overlapping features that are not well understood. Stressors that can induce cell death initiation include UV [26–29], ROS [30], and heat [31].

Cells have a number of intrinsic stress responses at their disposal to maintain homeostasis and cellular quality control. These include the catabolic degrative process of autophagy [17], oxidative stress response [18], heat shock response [19, 20], the Unfolded Protein Response (UPR) [21, 22] and DNA damage responses [23–25]. These protective responses are known to serve to counteract the effects of stress and preserve the cell. Do these same responses play any role when cells destined to die developmentally for proper homeostasis are exposed to stress? This problem is further compounded for cells that are specialized and of highly intricate structure such as neurons. The compartments of such morphologically complex cells have vastly different microenvironments which makes it plausible that their clearance mechanisms may be different.

How external stress and the cell fate of developmentally programmed cell elimination intersect is not well-studied. Outstanding questions include: Is preserving cellular integrity the only role of a stress response? Are there bonafide stress responses to permit programmed cell elimination under stress? Are canonical molecular players for cell removal and stress response involved for the removal of the different compartments of morphologically complex cells? We considered these questions and here address *in vivo* the question of how stress and stress response may impact the intrinsic fate of specialized cells destined to be eliminated and assess how cellular homeostasis may promote proper cell elimination under duress.

Previously, we have described a “tri-partite” developmental killing program for morphologically complex cells in the nematode *C. elegans* [3, 10]. In this program, Compartmentalized Cell Elimination, or CCE, three segments of the *C. elegans* tail-spike epithelial cell (TSC) (**FIG 1A-D**) and the sex-specific CEM neurons die in different ways [3, 10]. The TSC is a scaffolding epithelial cell that shapes the hyp10 tail-tip hypodermal cell (**FIG 1A**). During CCE, the TSC displays three degenerative morphologies – rounding soma, fragmenting proximal process, and retracting distal process (**FIG 1B**). The proximal process is the first to be fully removed (**FIG 1C**), leaving behind the soma and distal process remnants, which are cleared stochastically by different neighboring phagocytes (**FIG 1D**), with hyp10 acting as the process phagocyte, and an unidentified cell as the soma phagocyte. Following a forward genetic screen utilizing this system, we have previously molecularly characterized phagosome sealing, a poorly described yet critical step of phagocytosis, showing a requirement for the cell fusogen EFF-1 [10].

**Figure 1.**
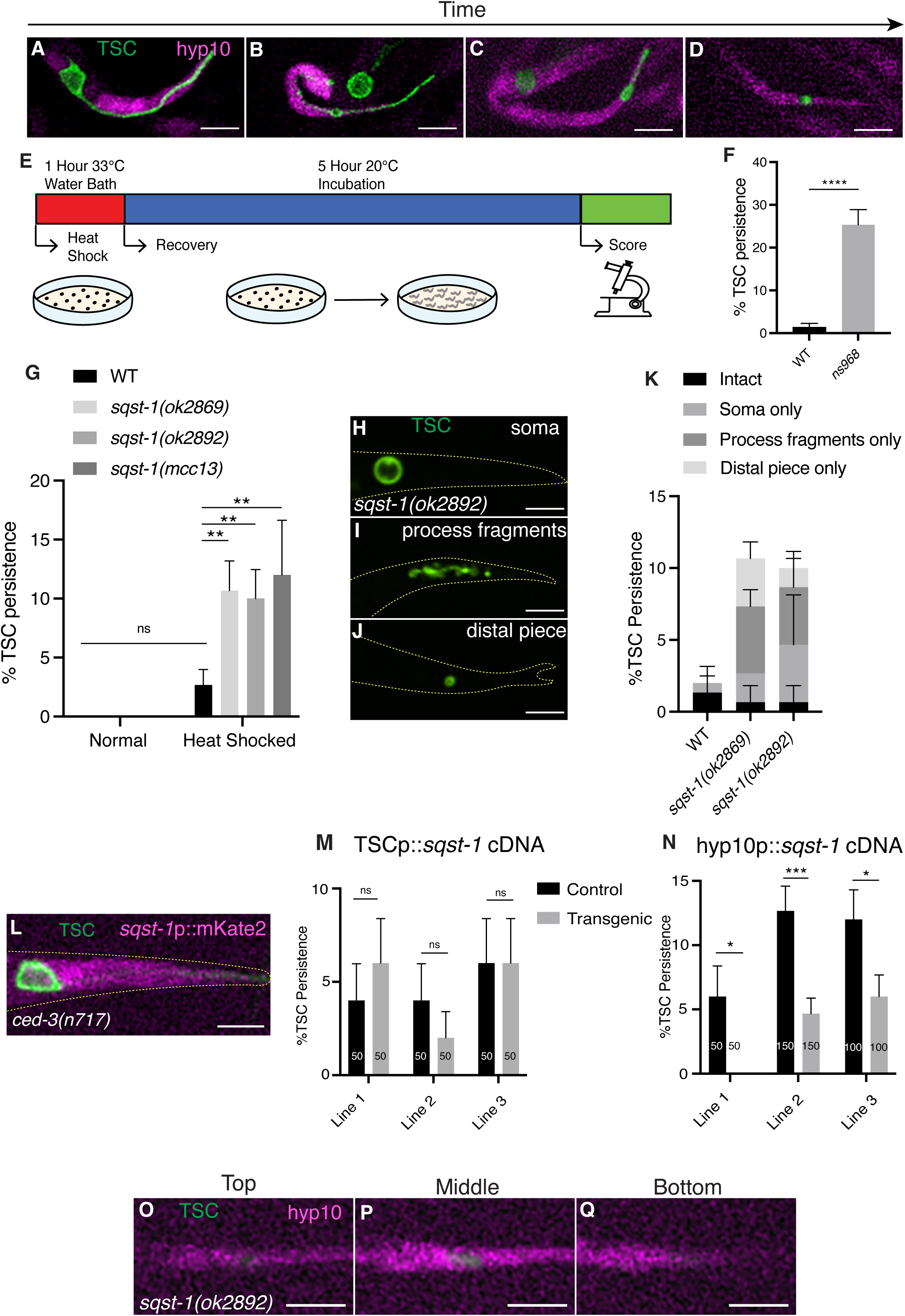
SQST-1/p62 functions in the hyp10 phagocyte to promote CCE following stress after TSC internalization (A-D) CCE of tail-spike cell (TSC, green) showing hyp10 cell (magenta) adjacent to TSC process**; (A)** Intact TSC. **(B)** Rounded TSC soma, fragmenting proximal process, retracting distal process. **(C)** Soma-distal remnants. **(D)** hyp10 engulfing the TSC process remnant. **(E)** Schematic of heat stress protocol. **(F)** Quantification of *ns968* mutant CCE defects. **(G)** Quantification of TSC persistence in wild-type vs *sqst-1(-).* **(H-J)** *sqst-1(ok2892)* mutant CCE defects following heat stress. **(K)** Quantification of *sqst-1(-)* phenotype categories. **(L)** *sqst-1* expression in *ced-3(n717)* mutant L1 larvae, with intact TSC. **(M)** Failure of TSC-specific rescue of *sqst-1(ok2892)* defect. **(N)** hyp10-specific rescue of *sqst-1(ok2892)* CCE defect. **(O-Q)** Persisting TSC remnant internalized by hyp10 phagocyte in *sqst-1(ok2892)* mutant at the top, middle, and bottom planes of hyp10. ns (not significant) *p* > 0.05, * *p* ≤ 0.05, ** *p* ≤ 0.01, *** *p* ≤ 0.001, **** *p* ≤ 0.0001.

We employed our CCE system, specifically the TSC, to address the questions above and provide here additional insight into the process of phagocytosis in the dual context of development and stress. We report additional lysosomal roles for stress-related genes to ensure efficient corpse processing and resolution in the phagosome, specifically under conditions of stress. We show that, following heat stress, the selective autophagy receptor SQST-1/p62 acts within the phagocyte to stabilize the oxidative stress transcription factor SKN-1/Nrf by negatively regulating a functional analog of KEAP1, WDR-23. Mammalian Nrf proteins are key mediators of various cytoprotective responses, with the Nrf2 responses to oxidative stress the best documented [32–34]. In C. *elegans,* SKN-1/Nrf plays roles in early development [35, 36] and also mediates conserved stress defense and detoxification responses and promotes longevity post-embryonically [37]. Our work shows that, in the context of CCE, SKN-1/Nrf stabilization promotes the transcription of the lysosomal trafficking associated gene *lyst-1/*LYST, validating, *in vivo,* prior genomics studies implicating an association [38]. This regulation appears to be important for the transition from the mature phagosome to phagolysosome stage at the level of lysosomal regulation. Our study highlights phagosome maturation during cell clearance as a new context for SKN-1/Nrf function, provides insights on roles of LYST-1/LYST, a poorly characterized protein, and highlights lysosome regulation as an important control point to ensure the efficient removal of developmentally killed cells under stress.

## Results

### Phagocytic SQST-1/p62 promotes CCE under stress cell non-autonomously

To test the hypothesis that developmental cell death is impacted by external stress, we examined whether CCE is achieved even under stress conditions and subjected our previously employed tail-spike cell membrane-targeted GFP (TSCp::myrGFP) transgenic animals to a heat stress paradigm (**FIG 1E**). Contrary to our hypothesis, under normal conditions, wild type animals under stress did not show significant TSC persistence (**FIG 1G**). We then considered that CCE may be accomplished even under stress via a stress response that promotes intrinsic cell elimination. We therefore considered a role for cellular quality control genes. We serendipitously came upon a relevant gene following a parallel forward genetic screen with the same transgenics under normal environmental conditions. We obtained a mutant from this screen, *ns968*, with remnants of both the TSC soma and process in the first larval (L1) stage, long after the cell should be cleared (**FIG 1F**). Following Whole Genome Sequencing, we noted a change in the gene *sqst-1*, which encodes the homolog of mammalian p62 sequestosome that results in a S350N change in exon 2.

p62 is a selective autophagy receptor which helps transport ubiquitinated proteins to the growing autophagosome [39–41] for their ultimate degradation. Autophagy involves the encompassing of a degradation target by an autophagosome vesicle to which lysosomes fuse, allowing for digestion by lysosomal hydrolases and destruction of the contents of the autophagosome [17]. Selective autophagy targets damaged organelles, invading pathogens and aggregated or unwanted proteins [41].

Intrigued, we attempted to confirm gene identity and tested three additional *sqst-1* alleles, two deletion alleles and a CRISPR/Cas9-engineered allele with the same lesion as in our original mutant. However, surprisingly, none of these showed the CCE defect of the screen mutant under normal conditions (**FIG 1G**). We then reasoned that the impact of a stress response gene mutation may only be realized under conditions of stress. As such, we subjected our *sqst-1*/p62 mutants to our heat-shock paradigm. Under these conditions, consistent with our idea, we observed robust CCE defects (**FIG 1G**), with a range of defects of both the soma and the process (**FIG 1H-K**), thus confirming gene identity. We then asked why our original mutant *ns968* displayed a CCE defect even in the absence of stress and postulated that this may be reconciled by the presence of an additional mutation. We elected to pursue work on the second mutation in the future and continued our present study with *sqst-1*/p62 mutants under stress conditions.

We first tested the expression of *sqst-1*/p62 using a transcriptional reporter for this gene (*sqst-1*p::mKate2). We observed expression in the hyp10 epithelial cell (**FIG 1L**), which we have previously shown serves both as the animal’s tail tip and as the phagocyte for the TSC process [10]. We moved forward with a focus on the TSC process and the hyp10 given our available tools. We next performed cell-specific rescue (**FIG 1M**) and found that introduction of *sqst-1*/p62 in the TSC did not rescue the mutant CCE heat shock defect. However, we observed rescue of the process phenotype by expression in hyp10 (**FIG 1N**). These data suggest that *sqst-1*/p62 functions in the hyp10 phagocyte to aid elimination of the TSC process cell non-autonomously. We next examined whether the TSC remnants following heat shock are internalized by hyp10 in *sqst-1*/p62 mutants. We visualized the location of these TSC process remnants (mKate2-PH) relative to hyp10 (iBlueberry) (**FIG 1O-Q**) and found that they were internalized. This suggests that the CCE defect following stress involves a step after corpse internalization, possibly during phagosome maturation and corpse processing. Mammalian p62 is known to bind directly to ubiquitinated targets via its UBA domain [42]. In worms, there is a single E1 ligase, UBA-1 [43]. When exposing *uba-1* mutants to our heat shock paradigm, we observed CCE defects similar to those of our *sqst-1*/p62 mutants (**FIG S1A-D**). This suggests that SQST-1 may act in its canonical capacity in selective autophagy together with UBA-1 to promote CCE and that these genes act in the same genetic pathway.

### SQST-1/p62 promotes SKN-1/Nrf2 function in CCE under stress

We next probed for the specific degradation target of SQST-1/p62. Previous studies in mammalian systems [44] have shown that mammalian SQSTM1/p62 facilitates the degradation of KEAP1, an E3 ubiquitin ligase adaptor [45–47] under oxidative stress conditions. KEAP1 is known to promote the degradation of the transcription factor Nrf2 under homeostatic conditions [48, 49]. Under oxidative stress conditions, KEAP1 is slated for degradation with the help of SQSTM1/p62, thereby permitting Nrf2 to translocate to the nucleus to promote transcription of oxidative stress response genes [44]. Bearing this model in mind, we tested mutants for *skn-1*, which encodes the homolog for Nrf1 and Nrf2 in nematodes [37, 50] and predicted that these mutants would phenocopy *sqst-1*/p62 mutants following heat stress. Indeed, *skn-1* mutants showed similar CCE defects as those for *sqst-1*/p62 (**FIG 2A-E**). Moreover, we found *skn-1* to be expressed in the phagocytes using a transcriptional reporter (**FIG 2F**); and that *skn-1* functions non-autonomously in the hyp10 phagocyte based on cell-specific rescue experiments (**FIG 2G, H**). *skn-1*/Nrf2 mutant TSC remnants were also found to be internalized by hyp10 (**FIG 2I-K**). We also examined the localization of SKN-1 following heat shock and found that endogenous SKN-1::GFP localizes to the nucleus following heat shock and that this is prevented in *sqst-1*/p62 mutants (**FIG 2L-P**). A *skn-1; sqst-1* double mutant was also not additive, suggesting these genes may act in the same pathway (**FIG 2A**). This positions SQST-1 and selective autophagy upstream of SKN-1/Nrf. Nrf2 is a transcription factor that acts in response to oxidative stress [37, 50]. It is typically found in the cytoplasm bound to KEAP1, which inhibits Nrf2 under normal physiological conditions [48, 49]. Following exposure to oxidative stress, KEAP1 releases Nrf2, allowing it to translocate to the nucleus and transcribe genes involved in cytoprotection [44]. While there is no known direct homolog of KEAP1 in *C. elegans,* WDR-23 is thought to degrade SKN-1/Nrf2 [51]. We also overexpressed *wdr-23* in hyp10 in wild-type animals and found this to phenocopy the *skn-1*/Nrf2 and *sqst-1*/p62 mutants (**FIG 2Q**), suggesting WDR-23 may be the KEAP1-like direct degradation target of SQST-1/p62 that negatively regulates SKN-1/Nrf2 in the context of CCE.

**Figure 2.**
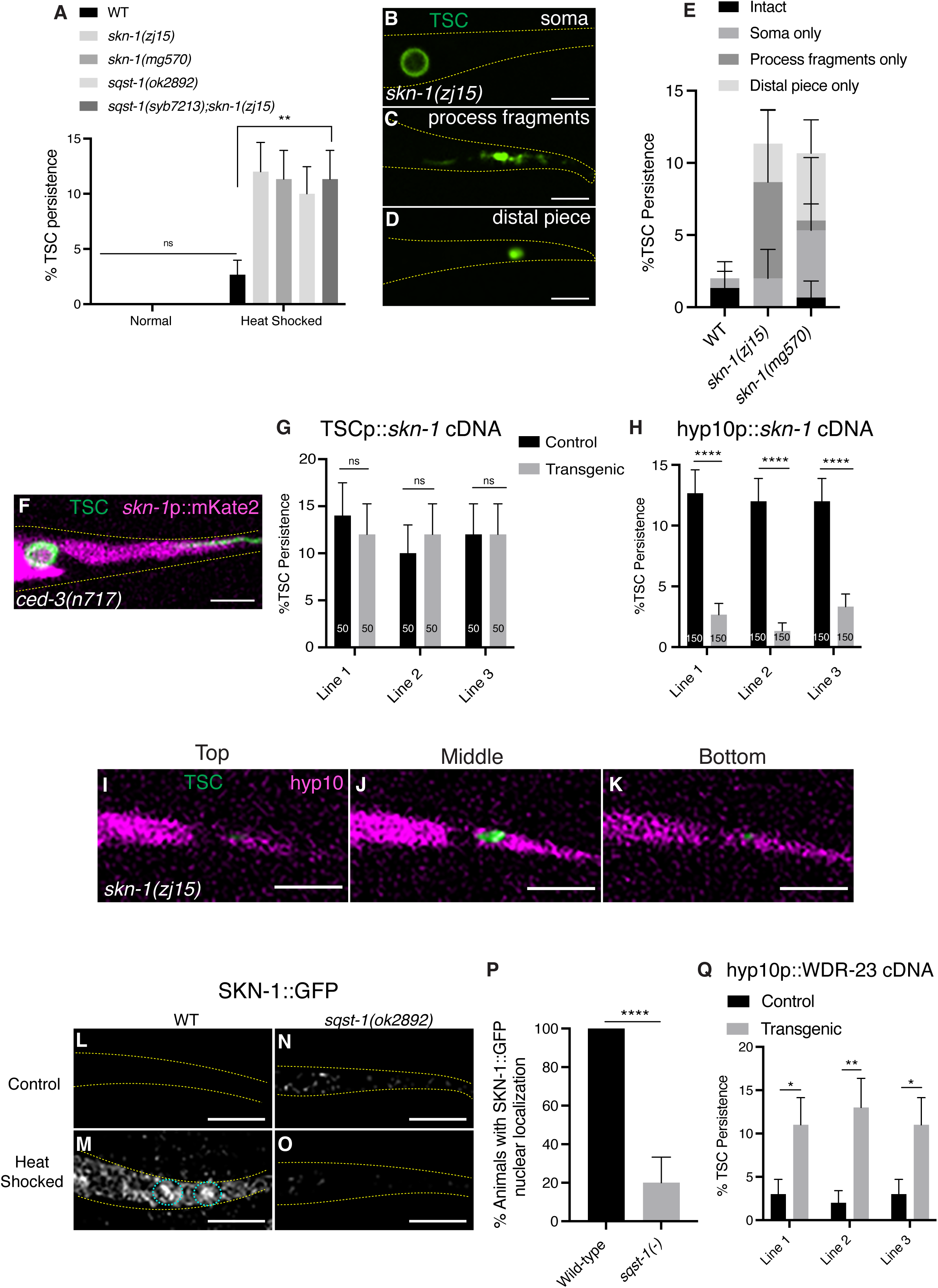
hyp10 phagocyte-specific SKN-1/Nrf2 promotes removal of internalized TSC remnants following stress in a SQST-1/p62 and WTS-1-dependent manner. **(A)** Quantification of TSC persistence in wild-type vs *skn-1(-).* (B-D) *skn-1(zj15)* mutant CCE defects following heat stress. **(E)** Quantification of *skn-1(-)* phenotype categories. (F) *skn-1* expression in *ced-3(n717)* mutant L1 larvae, with intact TSC. **(G)** Failure of TSC-specific rescue of *skn-1(zj15)* defect. **(H)** hyp10-specific rescue of *skn-1(zj15)* defect. **(I-K)** Persisting TSC remnant internalized by hyp10 phagocyte in *skn-1(zj15)* mutant at the top, middle, and bottom planes of hyp10. **(L-O)** Control and heat stress treated localization of endogenous SKN-1::GFP in wild-type and *sqst-1(ok2892)* mutants. **(P)** Quantification of (M, O). **(Q)** hyp10 driven overexpression of WDR-23 in wild-type animals. ns (not significant) *p* > 0.05, * *p* ≤ 0.05, ** *p* ≤ 0.01, *** *p* ≤ 0.001, **** *p* ≤ 0.0001.

### SKN-1/Nrf2 promotes *lyst-1* transcription in hyp10 phagocyte

We next sought to identify the transcriptional target of SKN-1/Nrf. Previous work has identified arrays of genes upregulated and downregulated in whole animals lacking *skn-1*/Nrf [38]. From this list of candidate SKN-1/Nrf targets, we tested *lyst-1*, which encodes the homolog of the lysosomal trafficking regulator LYST [52]. Mammalian LYST is a widely expressed gene encoding a protein that regulates intracellular protein trafficking from secretory lysosomes, but the mechanisms underlying its function are unknown [53–56]. The LYST gene is conserved and in *C. elegans lyst-1* is involved in gut granule formation and other lysosome-related organelle (LRO) biogenesis [52]. Aside from this, no roles of *lyst-1* in *C. elegans* have been described. As above, we subjected *lyst-1/*LYST mutants harboring TSCp::myrGFP to heat shock and observed CCE defects akin to those in *sqst-1*/p62 and *skn-1*/Nrf mutants, with *lyst-1; skn-1* double mutants not showing additive defects. (**FIG 3A-E)**. Our cell-specific rescue experiments suggest *lyst-1* also functions in the hyp10 phagocyte under stress conditions (**FIG 3F**). Additionally, we observed rescue of the *skn-1*/Nrf mutant phenotype when expressing *lyst-1* in hyp10 (**FIG 3G**) suggesting *lyst-1* functions downstream of SKN-1. We found that TSC fragments in *lyst-1* mutants are internalized by the hyp10 phagocyte as well (**FIG 3H-J**). Interestingly, we did not see *lyst-1* expression under basal conditions using a large *lyst-1* intron driving mKate2 (**FIG S2A**). However, following heat shock, *lyst-1* was expressed in the hyp10 phagocyte (**FIG S2B**). Moreover, loss of *sqst-1*/p62 or *skn-1*/Nrf2 prevented *lyst-1* expression in the hyp10 phagocyte even after heat shock (**FIG S2C-E**). We next mutated the consensus site of both the large last intron (atgacatt→ttgagatt) and promoter (attatcat→ttgagatt) of *lyst-1* using CRISPR/Cas9 gene editing on a strain in which GFP was incorporated into the endogenous locus of *lyst-1* (**FIG 3K**). While we did see *lyst-1* expression following heat shock in the wild-type background, fewer animals showed expression in the mutated strain (**FIG 3L-O**). These data support the idea that SKN-1/Nrf directly targets *lyst-1* regulatory regions to positively regulate *lyst-1* expression during stress.

**Figure 3.**
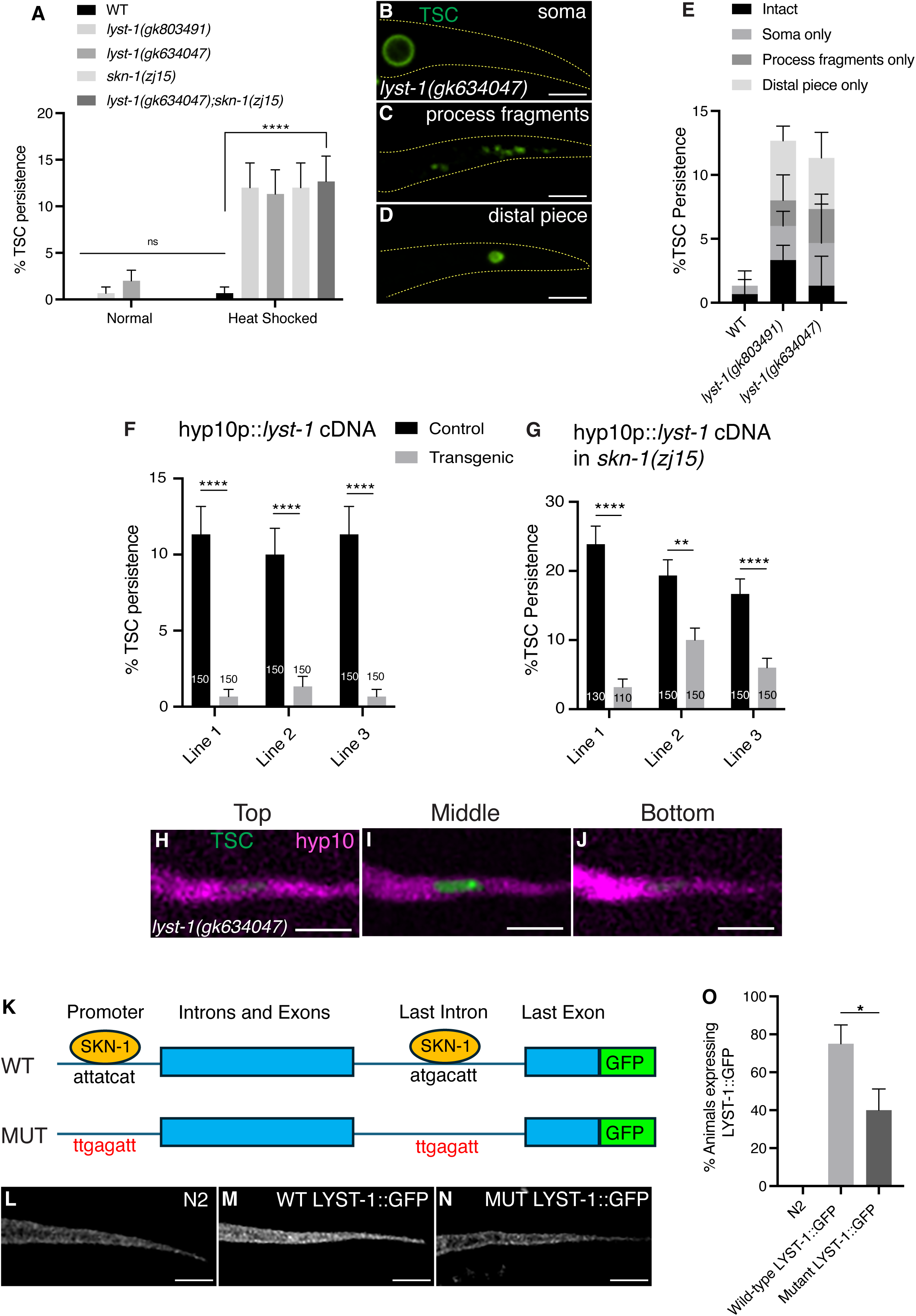
*lyst-1*/LYST promotes removal of the internalized TSC remnants following stress and is positively regulated transcriptionally by SKN-1/Nrf2 **(A)** Quantification of TSC persistence in wild-type vs *lyst-1(-).* (B-D) *lyst-1(gk634047)* mutant CCE defects following heat stress. **(E)** Quantification of *lyst-1(-)* phenotype categories. **(F)** hyp10-specific rescue of *lyst-1(gk634047)* defect. **(G)** hyp10-driven *lyst-1* rescue of *skn-1(zj15)* defect. **(H-J)** Persisting TSC remnant internalized by hyp10 phagocyte in *lyst-1 (gk634047)* mutant at the top, middle, and bottom planes of hyp10. **(K)** Simplified schematic of *lyst-1* gene structure showing mutated consensus promoter and last intron binding sites of SKN-1. **(L-N)** LYST-1 GFP binding site mutants. **(O)** Quantification of (L-N). ns (not significant) *p* > 0.05, * *p* ≤ 0.05, ** *p* ≤ 0.01, *** *p* ≤ 0.001, **** *p* ≤ 0.0001.

### The phagocytic stress response may affect phagolysosome formation

We next asked which step following corpse internalization is affected by this new SKN-1/Nrf dependent stress response, predicting that it may be the phagolysosome formation step, given the involvement of *lyst-1*/LYST. To evaluate/trace how far the mutant corpse has progressed inside the phagocyte during phagosome maturation in the different genetic backgrounds, we examined markers for various phagosome stages. Because of the putative function of *lyst-1/*LYST, we first tested a marker for RAB-7, which decorates late phagosomes that will subsequently fuse to incoming lysosomes [14, 15]. We found that in *sqst-1*/p62, *skn-1*/Nrf and *lyst-1* mutants, the TSC corpse is surrounded by RAB-7 signal (**FIG 4A-C’’**). This supports the notion that the corpse is arrested in the late phagosome prior to the membrane fusion of trafficked lysosomes. Consistent with this observation, we did not see enrichment with a phagocytic LAAT-1 marker for phagolysosomes (**FIG 4D-F’’**), which are formed only following the successful trafficking and fusion of lysosomes to the mature phagosome [57, 58]. We next examined mutants for *lmp-1*/Lamp-1 which encodes a protein important in lysosomes [59–62] under heat shock and observed that these mutants phenocopied *sqst-1*/p62, *skn-1*/Nrf and *lyst-1* mutants (**FIG 4G-K**). Moreover, we found *lmp-1* to also function in the hyp10 phagocyte based on cell-specific rescue experiments (**FIG 4L**).

**Figure 4.**
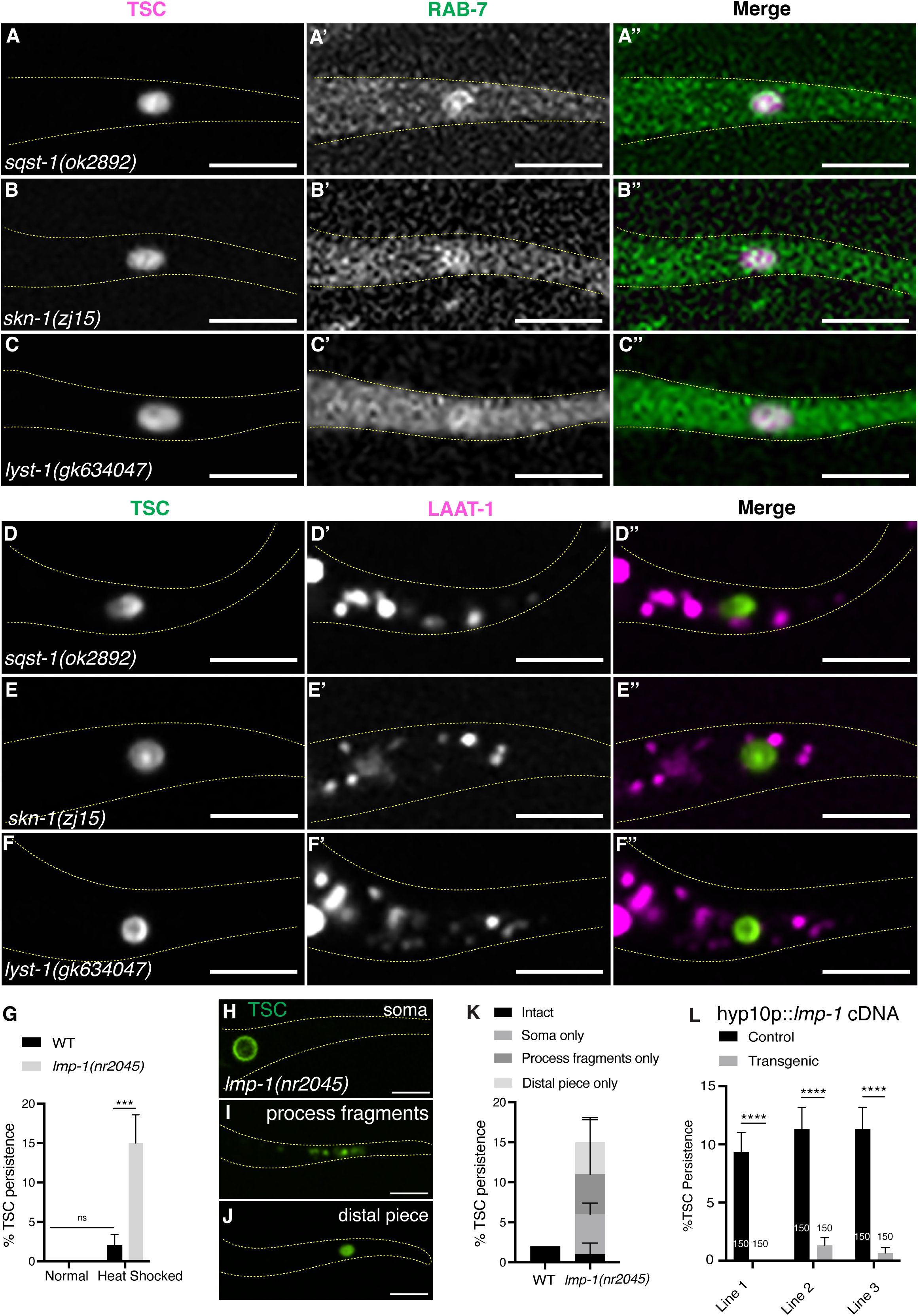
The SQST-1/SKN-1/LYST-1 stress response axis functions at the late-phagosome stage prior to phagolysosome formation (A-C’’) TSC remnant (magenta) localization relative to hyp10 RAB-7 (green) in *sqst-1(ok2892)*, *skn-1(zj15),* and *lyst-1(gk634047)* mutants. **(D-F’’)** TSC remnant (green) localization relative to hyp10 LAAT-1 (magenta) in *sqst-1(ok2892)*, *skn-1(zj15),* and *lyst-1(gk634047)* mutants. **(G)** Quantification of TSC persistence in wild-type vs *lmp-1(nr2045).* **(H-J)** *lmp-1(nr2045)* mutant CCE defects following heat stress. **(K)** Quantification of *lmp-1(-)* phenotype categories. **(L)** hyp10-specific rescue of *lmp-1(nr2045)* defect. ns (not significant) *p* > 0.05, * *p* ≤ 0.05, ** *p* ≤ 0.01, *** *p* ≤ 0.001, **** *p* ≤ 0.0001.

Collectively these data demonstrate a novel phagocytic stress response that ensures successful removal of a developmentally killed cell (**FIG 5, model**). We introduce new roles for and interactions between multiple genes in this context. We show a previously undescribed role and target for SKN-1/Nrf as well as evidence suggesting *C. elegans* WDR-23 functions analogous to mammalian KEAP1. We also present, to our knowledge, the first evidence directly implicating *lyst-1*/LYST in phagocytosis as well as in stress response. Taken together, our study establishes a direct genetic link between developmental cell elimination and stress response in the context of specialized cell elimination, which may have important implications to both neurodevelopment and neurodegeneration as well as general homeostasis.

**Figure 5.**
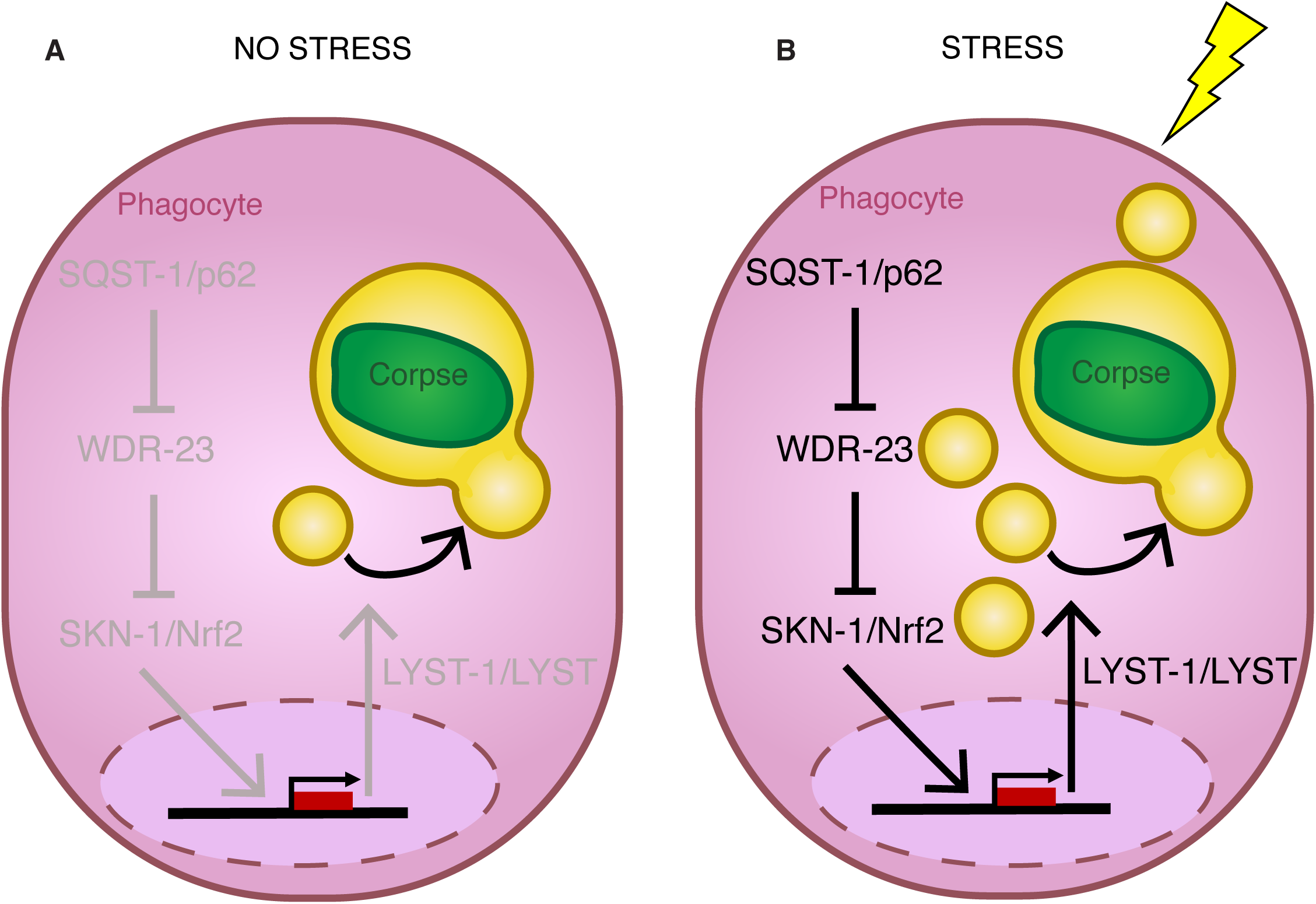
Model. SQST-1/p62 regulated SKN-1/Nrf promotes LYST-1/LYST function prior to phagolysosome formation for cell remnant resolution following stress. **(A)** Normal phagocytosis under non-stress conditions. **(B)** SQST-1/p62 promotes activity of SKN-1/Nrf2 to transcriptionally upregulate the lysosomal trafficking-associated gene *lyst-1* to ensure CCE during heat stress. This phagocytic stress response axis converges at lysosomes, following the late phagosome stage and prior to phagolysosome formation

## Discussion

The interplay between stress responses and development have been described in *C. elegans* in other contexts. For example, post-embryonic studies show that stress response genes are regulated by the extracellular matrix [63] and that somatic proteostasis and stress resilience are regulated in the reproductive system as a function of the status of the embryo [64]. In other experimental systems, prior studies have linked stress and phagocytosis specifically. For example, in the murine nervous system, it has been shown that stress hormones can induce synapse phagocytosis by astrocytes [65]. On the other hand, as shown in cultured cells, oxidative stress can inhibit the phagocytosis of apoptotic cells, despite phosphatidylserine (PS) externalization [66].

Here we have employed *C. elegans* Compartmentalized Cell Elimination to demonstrate a previously unreported link between developmentally programmed cell elimination and stress response at the point of the late step of phagosome maturation and define SKN-1/Nrf as a key molecular regulator. SKN-1/Nrf has been shown to have important roles in other developmental contexts, both embryonically and post-embryonically. During early *C. elegans* embryogenesis, SKN-1/Nrf specifies development of the endoderm and mesoderm [35, 36] with maternally-contributed SKN-1/Nrf functioning to establish the fate of the mesodermal precursor cell EMS [35, 36]. Post-embryonic SKN-1/Nrf is well known to mediate conserved stress defense and detoxification stress responses and promote longevity [37]. Mammalian Nrf proteins have been linked to cell removal. It has been shown that overexpression of Nrf1 sensitizes cells to apoptosis on serum depletion [67]. Nrf2 is upregulated in human renal tubule cells H_2_O_2_-mediated apoptotic injury [68]. Nrf2 has also been shown to promote macrophage function following bacterial infection [69]. Our study, by describing a new role of SKN-1/Nrf in phagocytosis of a developmentally killed cell, implicates Nrf proteins at large as versatile transcription factors that assure cells achieve their intended cell removal fate.

By validating prior genomics work [38] that SKN-1/Nrf targets *lyst-1*/LYST *in vivo,* our study also implicates phagolysosome formation as an important point of regulation for corpse resolution following stress for a developmentally killed cell. As mentioned, LYST is a conserved protein described as important for regulation of membrane dynamics and intracellular trafficking of lysosomes and lysosome related organelles (LROs) [56]. Prior work in the nematode model describes *lyst-1*/LYST’s involvement in gut granule and other LRO biogenesis, and that *lyst-1* mutants show decreased lysosome size [52]. Otherwise, this protein remains poorly characterized. Mutations in the human LYST gene have been implicated in different diseases, most notably the autosomal recessive immunodeficiency disease Chediak-Higashi syndrome [56, 70–79], marked by enlarged lysosomes and LROs [80]. An *in vivo* disease model does exist in the form of the *Lyst*-mutant ‘beige’ mouse [81]. Interestingly, in an immune context, work on macrophages suggests that *Lyst* mutations impair trafficking of the phagolysosome during the removal of bacteria [82], and LYST is important for phagosome maturation as it is required for recruitment of Rab7 during late stage endolysosomal maturation [83]. However, direct links of LYST proteins to cell corpse phagocytosis and development have not been made. In our model, we present an expanded role for a LYST protein in *C. elegans* in the regulation of phagocytosis as part of a developmental cell removal program potentiated by a stress response mediated by a SKN-1/Nrf. Further characterization of LYSTs and Nrfs in the context of cell corpse/debris resolution holds promise in the treatment of developmental diseases involving defects in cell elimination, as well as for other diseases marked by abnormal lysosome size and quantity.

Our identification of *lyst-1*/LYST as an effector of a phagocytic stress response during development underscores the importance of lysosomes to efficient cell corpse processing under stress. The fusion of lysosomes to phagosomes to form the phagolysosomes is a well-established essential step of phagosome maturation following the internalization of cargo [84]. Cargo could be in the form of various types of particles, such as microorganisms but also cells killed developmentally as in apoptosis with the goal of maintaining cellular homeostasis. Notably, lysosomes are established as efficient sensors of environmental changes and mediators of appropriate cellular adaptations to maintain homeostasis [85]. Our work highlights how regulation of these stress sensor organelles, or their association to the mature phagosome, is a critical point of regulation for programmed cell elimination, and that this regulation can be mediated by a bonafide stress response.

Our work also implicates WDR-23 in a new setting and identifies new regulation. In humans, Keap1 (Kelch-like ECH-associated protein 1) is the most highly studied regulator of NRF2, negatively regulating its protein levels and activity [48, 49]. No direct homolog for Keap1 is known in *C. elegans* [51] Instead, WDR-23, is a tryptophan aspartic acid WD40-repeat protein, has been shown to control SKN-1/Nrf [86]. Keap1 and WDR-23 are not similar in structure and their similarities are only at the mechanistic level. Nonetheless, the human genome has retained the WDR-23 homolog WDR23, despite the presence of Keap1. It has been shown that WDR23 can act as an alternative mode of regulation for NRF2 to Keap1 [51], for example in the nervous system [87]. This has been proposed to be highly relevant to cancer biology as cancer therapy is often associated with impairments in Keap1-dependent regulation of NRF2 [88]. Here we further validate WDR-23 as a functional analog to Keap1 in the regulation of SKN-1/Nrf in the new context of phagosome maturation, and additionally implicate p62 as a negative regulator. Interestingly, while p62 has been shown to negatively regulate Keap1 [44], an involvement in WDR23 regulation has not yet been demonstrated. Our finding together with the identification of additional roles and regulators of WDR23 could thus be of therapeutic value.

Our study also brings to light several intriguing questions. In this cell-cell interaction, both cells are subject to the same stress. The need to bolster the contribution of phagocyte lysosomes to the corpse resolution process may be owning to fundamental changes in the nature of the dying cell and resulting corpse (perhaps through its own stress adaptation) causing it to be difficult to process and digest. Given that the corpse is successfully internalized, there is presumably no change in membrane surface recognition and the thus far unidentified ‘eat-me’/recognition signal. Interestingly, in the case of phagocytosis during infection, some microorganisms can survive and grow within phagosomes. It has been shown that macrophages can expand their surface area to maintain membrane integrity to destroy such microorganisms. They can do so by inducing lysosome biogenesis, recruitment and insertion [89]. In the context of CCE, the stress-exposed dying/dead cell may need such additional mechanisms in place to be cleared and it will be interesting to learn what aspect lysosomal biology the SQST-1/SKN-1/LYST-1 axis regulates.

Our study identifies a phagocytic stress response. It will be interesting to tease apart whether the phagocytic response is due to the perception of external stress of the unideal yet internalized corpse. It is also possible that another phagosome maturation step is weakened under stress and LYST-1 function must compensate. There is also a temporal consideration as the phagocyte may be primed ahead of time by the environmental stress and then subsequently mounts its response to the corpse.

This study examined heat stress specifically. Follow-up work will examine whether other stressors have similar effects and whether specific forms of stress are relevant to developmental cell death, whether clearance following other types of cell death such as apoptosis or Linker cell-type death [3, 9] are also similarly impacted by stress and whether other steps of phagocytosis are regulated via stress response. Future screens combining the powerful CCE system with stress conditions may shed light on these and other open questions stemming from this study.

## Materials and Methods

### *C. elegans* methods

*C. elegans* strains were cultured using standard methods on *E. coli* OP50 and grown at 20°C. Wild-type animals were the Bristol N2 subspecies. For most TSC experiments, one of two integrated reporters were used: *nsIs435* or *nsIs685.* For hyp10 experiments, *nsIs836* was used. Integration of extrachromosomal arrays was performed using UV and trioxsalen (T2137, Sigma). Animals were scored at 20°C.

### Heat shock assay

For most experiments, embryos were subjected to heat shock in a water bath at 33°C for one hour and scored at the L1 stage following a recovery at 20°C for five hours, with the exception of *lyst-1* expression experiments, for which animals were subjected to heat shock at 37°C for one hour and scored immediately after.

### Imaging

Images were collected on a Nikon TI2-E Inverted microscope using a CFI60 Plan Apochromat Lambda 60x Oil Immersion Objective Lens, N.A. 1.4 (Nikon) and a Yokogawa W1 Dual Cam Spinning Disk Confocal. Images were acquired using NIS-Elements Advanced Research Package. For still embryo imaging, embryos were anesthetized using 0.5 M sodium azide. Larvae were paralyzed with 10mM sodium azide.

### Quantifications

For CCE defects, TSC death defects were scored at the L1 stage. Animals were mounted on slides on 2% agarose-M9 pads, paralyzed with 10mM sodium azide, and examined on a Zeiss Axio-Scope A1. The persisting TSC was identified by fluorescence based on its location and morphology. For SKN-1::GFP, animals were scored as presence or absence of nuclear GFP signal in the tail-tip at the L1 stage following heat shock. For LYST-1::GFP, animals were scored as presence or absence of GFP signal in the tail-tip at the L1 stage following heat shock.

### Worm strains used in this study

LGIV-*sqst-1(ok2869), sqst-1(ok2892)*, *sqst-1(mcc13), skn-1(zj15), skn-1(mg570), ced-3(n717), uba-1(it129)*

LGX-*lyst-1(gk634047), lyst-1(gk803491)*, *lyst-1(syb8801), lyst-1(syb8801 syb9206 syb9268), lmp-1(nr2045)*

### Plasmids and Transgenics

Plasmids were generated via Gibson cloning. Primer sequences and information on the construction of plasmids used in this study are provided in (Supplemental Table 1). The full list of transgenes is described in (Supplemental Table 2). The full length or fragment of the *aff-1* promoter was used to label the TSC. The *eff-1* promoter was used to label hyp10.

### CRISPR Cas9 genome editing

The alleles of *sqst-1(mcc13)* were made with a mutation resulting in a S350N change in exon 2 (the same as site as *ns968*). Mutants were generated using a co-injection strategy [90]. Guide crRNA, repair single-stranded DNA oligos, tracrRNA, and buffers were ordered from IDT. Guide crRNA used to generate *sqst-1(mcc13)* was 5’gatcattgaacgctcgacca-3’. The allele of *lyst-1*, syb8801[lyst-1::GFP] syb9206[lyst-1 last intron, (A11985bpT,C11989bpG] syb9268[lyst-1 promoter, g.-133_-139attatca˃ ttgagat] was generated by Suny Biotech (Suzhou, Jiangsu, China 215028).

### Statistics

Sample sizes and statistics were based on previous studies of CCE and the TSC [10]. Independent transgenic lines were treated as independent experiments. An unpaired two-tailed *t-*test was used for all persisting TSC quantifications (GraphPad Prism). For all figures, mean ± standard error of the mean (s.e.m.) is represented.

## Acknowledgments

AE and PG designed the experiments and wrote the manuscript. AE and AW performed the experiments. We thank Ginger Clark and Idara Ekong for technical assistance. We thank members of the Ghose lab for comments on the manuscript. We thank SUNY Biotech and Greg Hermann for strains. Some strains were provided by the CGC, which is funded by NIH Office of Research Infrastructure Programs (P40 OD010440).

## Funding

PG is a Cancer Prevention Research Institute of Texas (CPRIT) Scholar in Cancer Research (RR100091) and is also funded by a National Institutes of Health-National Institute of General Medical Sciences Maximizing Investigators’ Research Award (MIRA) (R35GM142489).

## Competing interests

The authors declare no competing interests.

## Figure Legends

**Supplement Figure 1. *uba-1*/UBA1 mutants show similar CCE defects to *sqst-1*/p62 mutants following stress. (A-C)** *uba-1(it129)* mutant CCE defects following heat stress. **(D)** Quantification of TSC persistence in wild-type vs *uba-1(it129).* ns (not significant) *p* > 0.05, * *p* ≤ 0.05, ** *p* ≤ 0.01, *** *p* ≤ 0.001, **** *p* ≤ 0.0001.

**Supplement Figure 2. *lyst-1*/LYST is expressed in hyp10 following stress and that expression is reduced in *sqst-1*/p62 and *skn-1*/Nrf mutants. (A)** Wild-type animal showing no expression of *lyst-* 1/LYST in hyp10 under basal conditions. **(B)** Wild-type animal showing expression of *lyst-1*/LYST in hyp10 following heat stress. (C) *sqst-1(ok2892)* mutant showing no expression of *lyst-1*/LYST in hyp10 following heat stress. (D) *skn-1(zj15)* mutant showing no expression of *lyst-1*/LYST in hyp10 following heat stress. **(E)** Quantification of B-D. ns (not significant) *p* > 0.05, * *p* ≤ 0.05, ** *p* ≤ 0.01, *** *p* ≤ 0.001, **** *p* ≤ 0.0001.

**Supplement Table 1. Plasmids used in this study**

**Supplement Table 2. List of transgenes and strains**

**Supplement Movie 1. Expression of *sqst-1*/p62 in hyp10**

**Supplement Movie 2. *sqst-1(ok2892)* mutant remnants appear internalized by hyp10**

**Supplement Movie 3. Expression of *skn-1*/Nrf in hyp10**

**Supplement Movie 4. *skn-1(zj15)* mutant remnants appear internalized by hyp10**

**Supplement Movie 5. SKN-1/Nrf does not localize to the nuclei of hyp10 under basal conditions in wild-type**

**Supplement Movie 6. SKN-1/Nrf localizes to the nuclei of hyp10 following heat stress in wild-type**

**Supplement Movie 7. SKN-1/Nrf does not localize to the nuclei of hyp10 under basal conditions in *sqst-1(ok2892)* mutants**

**Supplement Movie 8. SKN-1/Nrf does not localize to the nuclei of hyp10 following heat stress in *sqst-1(ok2892)* mutants**

**Supplement Movie 9. *lyst-1(gk634047)* mutant remnants appear internalized by hyp10**

**Supplement Movie 10. N2 animal following heat stress**

**Supplement Movie 11. Wild-type animal expressing LYST-1::GFP following heat stress**

**Supplement Movie 12. LYST-1::GFP binding site mutant following heat stress**

**Supplement Movie 13. RAB-7::GFP localization around remnant of *sqst-1(ok2892)* mutant**

**Supplement Movie 14. RAB-7::GFP localization around remnant of *skn-1(zj15)* mutant**

**Supplement Movie 15. RAB-7::GFP localization around remnant of *lyst-1(gk634047)* mutant**

**Supplement Movie 16. LAAT-1::mCherry showing lysosomes unfused to remnant of *sqst-1(ok2892)* mutant**

**Supplement Movie 17. LAAT-1::mCherry showing lysosomes unfused to remnant of *skn-1(zj15)* mutant**

**Supplement Movie 18. LAAT-1::mCherry showing lysosomes unfused to remnant of *lyst-1(gk634047)* mutant**

**Supplement Movie 19. Wild-type animal showing no expression of *lyst-1* intron-driven mKate2 under basal conditions**

**Supplement Movie 20. Wild-type animal showing expression of *lyst-1* intron-driven mKate2 following heat stress**

**Supplement Movie 21. *sqst-1(ok2892)* mutant not showing expression of *lyst-1* intron-driven mKate2 following heat stress**

**Supplement Movie 22. *skn-1(zj15)* mutant not showing expression of *lyst-1* intron-driven mKate2 following heat stress**

## Notes

### Competing Interest Statement

The authors have declared no competing interest.

