## Supplementary figures and images for "SQST-1/p62-regulated SKN-1/Nrf mediates a phagocytic stress response via transcriptional activation of *lyst-1*/LYST"

### Supplemental Figure 1

**Elkhalil et. al, SUPPLEMENT FIGURE 1**

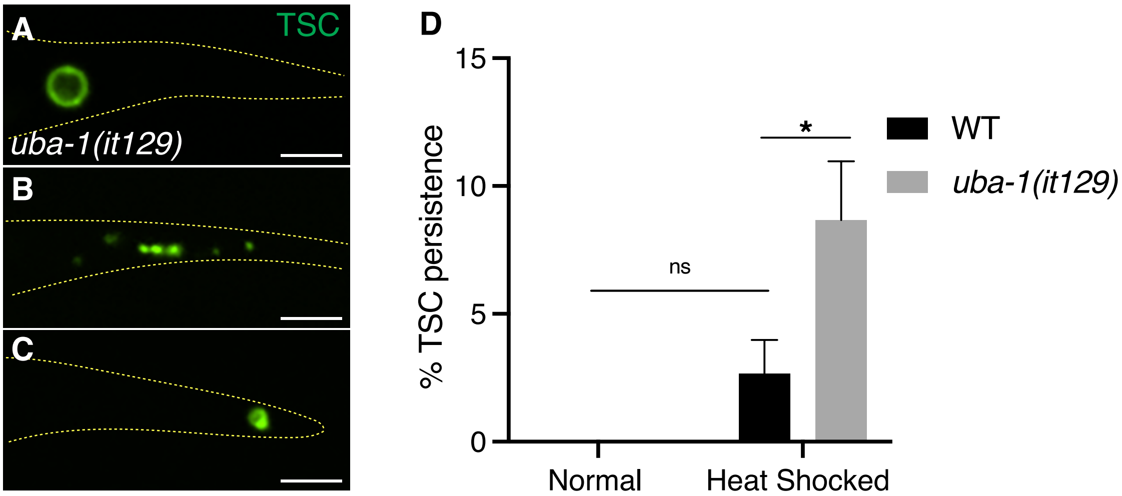

### Supplemental Figure 2

**Elkhalil et. al, SUPPLEMENT FIGURE 2**

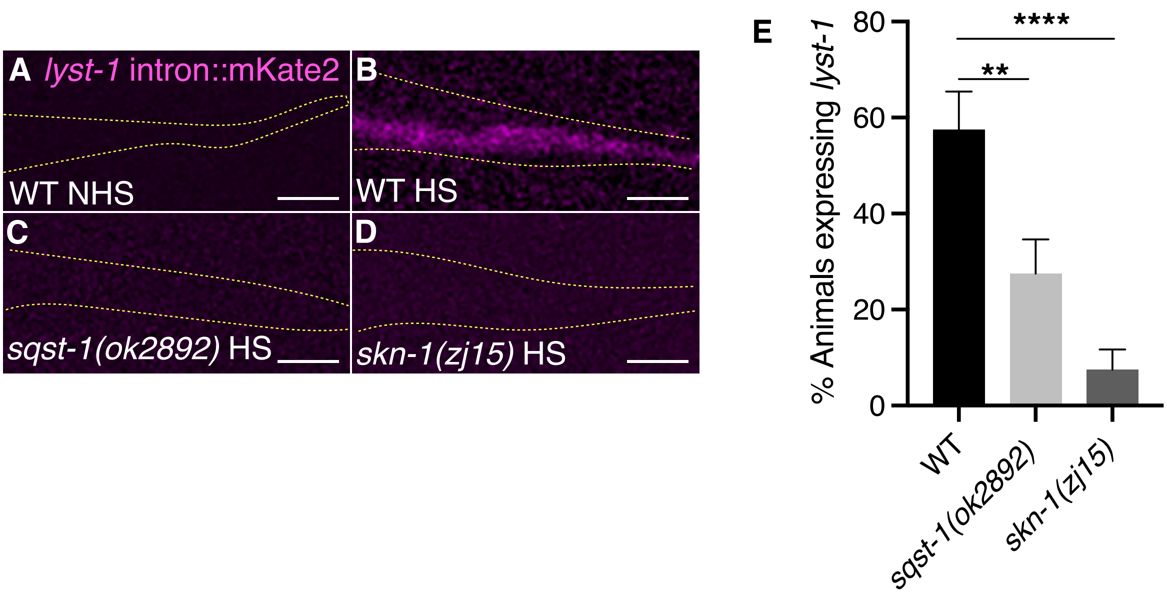
