## Supplemental Table 1 for "SQST-1/p62-regulated SKN-1/Nrf mediates a phagocytic stress response via transcriptional activation of *lyst-1*/LYST"

**Supplement Table 1: Plasmids used in this study**

| Plasmid number | Primers used | Sequences |
|---|---|---|
| pPG302 (*sqst-1* expression) | oAE15 mKate2 Vector backbone_F | ATGGTCTCCGAGCTCATTAAcGAAAAC |
| | oAE16 mKate2 Vector backbone_R | GATCCTCTAGAGTCGACCTGCAGGC |
| | oAE17 *sqst-1* pro insert-F | GAAATAAGCTTGCATGCCTGCAGGTCGACTCTAGAGGATCagatcattcaaagatgaaatccattagatt |
| | oAE18 *sqst-1* pro insert-R | GCTTCATATGCATGTTTTCgTTAATGAGCTCGGAGACCATctgagtaaaatgagaagttatttgaaagtg |
| pPG312 (*skn-1* expression) | oAE15 mKate2 Vector backbone_F | ATGGTCTCCGAGCTCATTAAcGAAAAC |
| | oAE16 mKate2 Vector backbone_R | GATCCTCTAGAGTCGACCTGCAGGC |
| | oAE31 *skn-1* pro insert_F | GAAATAAGCTTGCATGCCTGCAGGTCGACTCTAGAGGATCtgctcacagatctcaaagctgcgtgtg |
| | oAE32 *skn-1* pro insert_R | GCTTCATATGCATGTTTTCgTTAATGAGCTCGGAGACCATctgaaaatttggaattatttttgggaatatcg |
| pPG313 (*sqst-1* tsc rescue) | oYM14 *sqst-1* gDNA insert_F | tccatactttctcatttcataatattcggaacGGCGCGCCaaaATGGCTGCTGCATCATCCGCTCC |
| | oYM15 *sqst-1* gDNA insert_R | tacctttgggtcctttggccaatcccggggatcctctagaTTAGTGAAGAAGCGCCTGAAGACACATC |
| | oKJ43 TSCpro backbone_F | GGCGCGCCgttccgaatattatg |
| | oKJ44 TSCpro backbone_R | GAATTCcaactgagcgccggtc |
| pPG325 (*skn-1* tsc rescue) | oYM37 TSCpro backbone_F | tctagaggatccccgggattggcc |

| | | |
|---|---|---|
| | oYM38 TSCpro backbone_R | GGCGCGCCgttccgaatattatgaaa |
| | oYM39 *skn-1a* cDNA insert_F | tccatactttctcatttcataatattcggaacGGCGCGCCaaaATGGGCGGTTCATCACGCCGTCA |
| | oYM40 *skn-1a* cDNA insert_R | tacctttgggtcctttggccaatcccggggatcctctagaTCAGATGTAATGGGACATCTTGTCGTG |
| pPG336 (*lyst-1* intron expression) | oAE40 mKate2 Vector backbone_F | ATGGTCTCCGAGCTCATTAAcGAAAAC |
| | oAE41 mKate2 Vector backbone_R | GATCCTCTAGAGTCGACCTGCAGGCATG |
| | oAE44 *lyst-1* intron insert_F | GAAATAAGCTTGCATGCCTGCAGGTCGACTCTAGAGGATCtgctttttatatttcgttttaatgttttgcag |
| | oAE45 *lyst-1* intron insert_R | GCTTCATATGCATGTTTTCgTTAATGAGCTCGGAGACCATctgaaaaaataataactttttaaaccctaac |
| pPG359 (*sqst-1* hyp10 rescue) | oAE70 vector backbone_F | GGCCGGCCcagtcagtgcggccgc |
| | oAE71 vector backbone_R | gctgtctcatcctactttcacctagttaac |
| | oAE72 hyp10pro insert_F | cgccaagcttgcatgcgcggccgcactgactgGGCCGGCCttttgatactttttaatcaaaaagtttaccgc |
| | oAE73 hyp10pro insert_R | GAACTTGCATTTGGTGAGGAGCGGATGATGCAGCAGCCATttttaaaacaaaaaaagatgccttcctattg |
| | oAE74 *sqst-1* gDNA insert_F | gttagtttccaataggaaggcatctttttttgttttaaaaATGGCTGCTGCATCATCCGCTCCTCAC |
| | oAE75 *sqst-1* gDNA insert_R | aagacaagcagttaactaggtgaaagtaggatgagacagcTTAGTGAAGAAGCGCCTGAAGACACATC |
| pPG360 (*skn-1* hyp10 rescue) | oAE70 vector backbone_F | GGCCGGCCcagtcagtgcggccgc |

| | | |
|---|---|---|
| | oAE71 vector backbone_R | gctgtctcatcctactttcacctagttaac |
| | oAE76 hyp10pro backbone_F | cgccaagcttgcatgcgcggccgcactgactgGGCCGGCCttttgatactttttaatcaaaaagtttaccgc |
| | oAE77 hyp10pro backbone_R | CCGACGTACTTCGCTGACGGCGTGATGAACCGCCCATtttttttaaaacaaaaaaagatgccttcctattgg |
| | oAE78 *skn-1a* cDNA insert_F | gttagtttccaataggaaggcatctttttttgttttaaaaaaaATGGGCGGTTCATCACGCCGTCAGC |
| | oAE79 *skn-1a* cDNA insert_R | aagacaagcagttaactaggtgaaagtaggatgagacagcTCAGATGTAATGGGACATCTTGTCGTGAC |
| pPG370b (*lyst-1* hyp10 rescue) | oAE98 hyp10pro backbone_F | gctgtctcatcctactttcacctagtt |
| | oAE99 hyp10pro backbone_R | ttttaaaacaaaaaaagatgccttcctattg |
| | oAE114 *lyst-1* cDNA (1st half) insert_F | gttagtttccaataggaaggcatctttttttgttttaaaaaaaATGGAAAAGATCCGCTCACCATCACT |
| | oAE115 *lyst-1* cDNA (1st half) insert_R | TAGATTTAATTGGACGATTTTTGTTATCTTGTCCGAGTAAATGTTATGGTTAATAATAGCTTCATACATCTC |
| | oAE116 *lyst-1* cDNA (2nd half) insert_F | CCCTCATAGAGATGTATGAAGCTATTATTAACCATAACATTTACTCGGACAAGATAACAAAAATCGTCC |
| | oAE117 *lyst-1* cDNA (2nd half) insert_R | aagacaagcagttaactaggtgaaagtaggatgagacagcTCAAGTTCTTATTTTGAATCTCCACGTTTTG |
| pPG404 (*lmp-1* hyp10 rescue) | oAE138 hyp10pro backbone_F | gctgtctcatcctactttcacctagtt |
| | oAE139 hyp10pro backbone_R | ttttaaaacaaaaaaagatgccttcctattg |

| | | |
|---|---|---|
| | oAE140 *lmp-1* cDNA insert_F | gttagtttccaataggaaggcatctttttttgttttaaaaaaaATGTTGAAATCGTTTGTCATCTTGTTTG |
| | oAE141 *lmp-1* cDNA insert_R | aagacaagcagttaactaggtgaaagtaggatgagacagcTTAGACGCTGGCATATCCTTGTCTC |
| pPG422 (*wdr-23* Overexpression) | oAE163 hyp10pro backbone_F | ttttaaaacaaaaaaagatgccttcctattgg |
| | oAE164 hyp10pro backbone_R | gctgtctcatcctactttcacctagttaac |
| | oAE165 *wdr-23* cDNA insert_F | gttagtttccaataggaaggcatctttttttgttttaaaaaaaATGGGCAACTGGATAACGTCGACG |
| | oAE166 *wdr-23* cDNA insert_R | aagacaagcagttaactaggtgaaagtaggatgagacagcTTAATTTTGAGAGATGCTGCTCGATGAGC |
