## Supplemental Table 2 for "SQST-1/p62-regulated SKN-1/Nrf mediates a phagocytic stress response via transcriptional activation of *lyst-1*/LYST"

**Supplement Table 2: List of transgenes and strains**

| Strain | Genotype | Comments |
|---|---|---|
| TSC125 | T12G3.1(ok2869); *nsIs435* | T12G3.1=*sqst-1; nsIs435=aff-1*p::myrGFP |
| TSC126 | T12G3.1(ok2892); *nsIs435* | T12G3.1=*sqst-1; nsIs435=aff-1*p::myrGFP |
| TSC148 | *skn-1(zj15); nsIs435* | *nsIs435=aff-1*p::myrGFP |
| TSC149 | *skn-1(mg570); nsIs435* | *nsIs435=aff-1*p::myrGFP |
| TSC156 | *uba-1(it129); nsIs435* | *nsIs435=aff-1*p::myrGFP |
| TSC174 | *ced-3(n717); nsIs435; mccEx096* | *mccEx096*=pPG312; *nsIs435=aff-1*p::myrGFP |
| TSC175 | *sqst-1(ok2892); nsIs435; mccEx097* | *mccEx097*=pPG313; *nsIs435=aff-1*p::myrGFP |
| TSC176 | *sqst-1(ok2892); nsIs435; mccEx098* | *mccEx098*=pPG313; *nsIs435=aff-1*p::myrGFP |
| TSC177 | *sqst-1(ok2892); nsIs435; mccEx099* | *mccEx099*=pPG313; *nsIs435=aff-1*p::myrGFP |
| TSC193 | *ced-3(n717); nsIs435; mccEx110* | *mccEx110*=pPG302; *nsIs435=aff-1*p::myrGFP |
| TSC214 | *lyst-1(gk803491); nsIs435* | *nsIs435=aff-1*p::myrGFP |
| TSC215 | *lyst-1(gk634047); nsIs435* | *nsIs435=aff-1*p::myrGFP |
| TSC220 | *ced-3(n717); nsIs435; mccEx126* | *mccEx126*=pPG336; *nsIs435=aff-1*p::myrGFP |
| TSC221 | *ced-3(n717); nsIs435; mccEx127* | *mccEx127*=pPG336; *nsIs435=aff-1*p::myrGFP |
| TSC222 | *ced-3(n717); nsIs435; mccEx128* | *mccEx128*=pPG336; *nsIs435=aff-1*p::myrGFP |
| TSC223 | *skn-1(zj15); nsIs435; mccEx119* | *mccEx119*=pPG325; *nsIs435=aff-1*p::myrGFP |
| TSC224 | *skn-1(zj15); nsIs435; mccEx120* | *mccEx120*=pPG325; *nsIs435=aff-1*p::myrGFP |
| TSC225 | *skn-1(zj15); nsIs435; mccEx121* | *mccEx121*=pPG325; *nsIs435=aff-1*p::myrGFP |
| TSC232 | *lmp-1(nr2045); nsIs435* | *nsIs435=aff-1*p::myrGFP |
| TSC254 | *ced-3(n717); mccIs017* | *mccIs017*=pPG336 |
| TSC265 | *sqst-1(mcc13); nsIs435* | *mcc13=sqst-1(mcc13)*.1358 G -> A; *nsIs435=aff-1*p::myrGFP. |
| TSC279 | T12G3.1(ok2892); *lyst-1(gk634047); nsIs435* | T12G3.1=*sqst-1*; *nsIs435=aff-1*p::myrGFP |
| TSC280 | *skn-1(zj15); lyst-1(gk634047); nsIs435* | *nsIs435=aff-1*p::myrGFP |
| TSC281 | N2; *mccIs017; nsIs435* | *mccIs017*=pPG336 |
| TSC315 | T12G3.1(ok2892); *nsIs435; mccEx161* | *mccEx161*=pPG359; T12G3.1=*sqst-1*; *nsIs435=aff-1*p::myrGFP |
| TSC316 | T12G3.1(ok2892); *nsIs435; mccEx162* | *mccEx162*=pPG359; T12G3.1=*sqst-1*; *nsIs435=aff-1*p::myrGFP |

| | | |
|---|---|---|
| TSC317 | T12G3.1(ok2892); *nsIs435; mccEx163* | *mccEx163*=pPG359; T12G3.1=*sqst-1*; *nsIs435=aff-1*p::myrGFP |
| TSC318 | *skn-1(zj15); nsIs435; mccEx164* | *mccEx164*=pPG360; *nsIs435=aff-1*p::myrGFP |
| TSC319 | *skn-1(zj15); nsIs435; mccEx165* | *mccEx165*=pPG360; *nsIs435=aff-1*p::myrGFP |
| TSC320 | *skn-1(zj15); nsIs435; mccEx166* | *mccEx166*=pPG360; nsIs435=*aff-1*p::myrGFP |
| TSC324 | T12G3.1(ok2892); mccIs017; nsIs435 | *mccIs017*=pPG336; T12G3.1=*sqst-1*; *nsIs435=aff-1*p::myrGFP |
| TSC370 | *skn-1(zj15); mccIs017; nsIs435* | *mccIs017*=pPG336; *nsIs435=aff-1*p::myrGFP |
| TSC392 | *lyst-1(gk634047); nsIs435; mccEx210* | *mccEx210*=pPG370b; *nsIs435=aff-1*p::myrGFP |
| TSC393 | *lyst-1(gk634047); nsIs435; mccEx211* | *mccEx211*=pPG370b; *nsIs435=aff-1*p::myrGFP |
| TSC394 | *lyst-1(gk634047); nsIs435; mccEx212* | *mccEx212*=pPG370b; *nsIs435=aff-1*p::myrGFP |
| TSC429 | *nsIs685; nsIs836* | *nsIs685=aff-1*p::mKate2; *nsIs836=eff-1*p::iBlueberry |
| TSC430 | T12G3.1(ok2892); *nsIs685; nsIs836* | T12G3.1=*sqst-1*; *nsIs685=aff-1*p::mKate2; *nsIs836=eff-1*p::iBlueberry |
| TSC432 | *skn-1(zj15); nsIs685; nsIs836* | *nsIs685=aff-1*p::mKate2; *nsIs836=eff-1*p::iBlueberry |
| TSC473 | *lmp-1(nr2045); nsIs435; mccEx234* | *mccEx234*=pPG404 |
| TSC474 | *lmp-1(nr2045); nsIs435; mccEx235* | *mccEx235*=pPG404 |
| TSC475 | *lmp-1(nr2045); nsIs435; mccEx236* | *mccEx236*=pPG404 |
| TSC480 | T12G3.1(ok2892); *nsIs435; nsEx5971* | T12G3.1=*sqst-1*; *nsEx5971=ced-1*p::LAAT-1::mCherry; *nsIs435=aff-1*p::myrGFP |
| TSC481 | *skn-1(zj15); nsIs685; nsEx5975* | *nsEx5975=eff-1*p::GFP::RAB-7; *nsIs685=aff-1*p::mKate2 |
| TSC511 | *skn-1(zj15); nsIs435; mccEx241* | *mccEx241*=pPG370b; *nsIs435=aff-1*p::myrGFP |
| TSC512 | *skn-1(zj15); nsIs435; mccEx242* | *mccEx242*=pPG370b; *nsIs435=aff-1*p::myrGFP |
| TSC513 | *skn-1(zj15); nsIs435; mccEx243* | *mccEx243*=pPG370b; *nsIs435=aff-1*p::myrGFP |
| TSC585 | *nsIs435; mccEx247* | *mccEx247*=pPG422; *nsIs435=aff-1*p::myrGFP |
| TSC586 | *skn-1(zu169); geIs7; nsIs685; nsIs836* | *geIs7* [*skn-1b*::GFP]; *nsIs685=aff-1*p::mKate2; *nsIs836=eff-1*p::iBlueberry |

| | | |
|---|---|---|
| TSC587 | T12G3.1(ok2892); *geIs7; nsIs685; nsIs836* | T12G3.1=*sqst-1*; *geIs7* [*skn-1b*::GFP]; *nsIs685=aff-1*p::mKate2; *nsIs836=eff-1*p::iBlueberry |
| TSC589 | T12G3.1(ok2892); *nsIs685; nsEx5975* | T12G3.1=*sqst-1*; *nsEx5975=eff-1*p::GFP::RAB-7; *nsIs685=aff-1*p::mKate2 |
| TSC590 | *skn-1(zj15); nsIs435; nsEx5971* | *nsEx5971=ced-1*p::LAAT-1::mCherry; *nsIs435=aff-1*p::myrGFP |
| TSC591 | *lyst-1(gk634047); nsIs435; nsEx5971* | *nsEx5971=ced-1*p::LAAT-1::mCherry; *nsIs435=aff-1*p::myrGFP |
| TSC625 | *nsIs435; mccEx256* | *mccEx256*=pPG422; *nsIs435=aff-1*p::myrGFP |
| TSC626 | *nsIs435; mccEx257* | *mccEx257*=pPG422; *nsIs435=aff-1*p::myrGFP |
| TSC656 | *lyst-1(gk634047); mccIs099; nsEx5975* | *mccIs099=aff-1*p::myrmCherry; *nsEx5975=eff-1*p::GFP::RAB-7 |
| TSC669 | *lyst-1(gk634047); mccIs099; mccEx062* | *mccIs099=aff-1*p::myrmCherry; *mccEx062=eff-1*p(-4200 ->-4450)::GFP |
| PHX7213 | *sqst-1(syb7213) skn-1(zj15)* | |
| PHX8801 | *lyst-1(syb8801)* | |
| PHX9268 | *lyst-1(syb8801 syb9206 syb9268)* | |
| OS12750 | *ns968; nsIs435* | *ns968: sqst-1* S350N change in exon 2; *nsIs435=aff-1*p::myrGFP |
